# Integrating transcriptomic network reconstruction and QTL analyses reveals mechanistic connections between genomic architecture and *Brassica rapa* development

**DOI:** 10.1101/540740

**Authors:** Robert L. Baker, Wen Fung Leong, Marcus T. Brock, Matthew J. Rubin, R. J. Cody Markelz, Stephen Welch, Julin N. Maloof, Cynthia Weinig

## Abstract

Plant developmental dynamics can be heritable, genetically correlated with fitness and yield, and undergo selection. Therefore, characterizing the mechanistic connections between the genetic architecture governing plant development and the resulting ontogenetic dynamics of plants in field settings is critically important for agricultural production and evolutionary ecology. We use a hierarchical Bayesian Function-Valued Trait (FVT) approach to estimate *Brassica rapa* growth curves throughout ontogeny, across two treatments and in two growing seasons. We find that the shape of growth curves is relatively plastic across environments compared to final height, and that there are trade-offs between growth rate and duration. We determined that combining FVT Quantitative Trait Loci (QTL) and genes/eigengene expression identified via transcriptomic co-expression network reconstructions best characterized phenotypic variation. Further, targeted eQTL analyses identified regulatory hotspots that colocalized with FVT QTL and co-expression network identified genes and mechanistically link FVT QTL with structural trait variation throughout development in agroecologically relevant field settings.

## INTRODUCTION

Plant developmental genetics are correlated with fitness and yield (Baker *et al.* 2015; Kulbaba *et al.* 2017). Therefore, characterizing the mechanistic connections between the genetic architecture governing plant development and the resulting ontogenetic dynamics of plants in field settings is critically important to improving agricultural production and understanding evolutionary performance. Forward genetic approaches such as quantitative trait mapping are an attractive method of characterizing genetic architecture because they do not require *a priori* information such as candidate loci and can be used to describe pleiotropic and epistatic loci as well as polygenic traits (Prioul *et al.* 1997; Mackay 2013; Csilléry *et al.* 2018). Transcriptomic co-expression analyses and expression QTL (eQTL) have also been used to identify the underlying genetic architecture responsible for phenotypic variation (e.g. Nozue *et al.* 2018). Recently, combining information from genomic association studies and transcriptomic expression analyses has been used to pinpoint candidate genes (Hitzemann *et al.* 2003; Li *et al.* 2018; Luo *et al.* 2018; Schaefer *et al.* 2018). However, co-expression network analyses can also provide insight into the mechanistic connections between QTL genotypes and phenotypes. Here, we ask whether QTL, co-expression analyses, or a combination thereof best predict phenotypic variation. In combination with a targeted eQTL analyses in agroecologically relevant field settings, we characterize the mechanistic connections between the genomic architecture, transcriptomic expression networks, and phenotypic variation throughout plant development.

Development rarely occurs in discrete steps, yet developmental data are typically collected at multiple distinct but inter-dependent time points. Function-Valued Trait (FVT) modeling is one method of estimating the underlying continuous nature of development and avoiding complicated repeated measures analyses, which often compromise statistical power in downstream analyses (Wu *et al.* 1999; Griswold *et al.* 2008). One approach to FVT modeling involves fitting mathematical functions to discrete data to estimate continuous curves that represent the change of a trait or character as a function, typically of time (Kingsolver *et al.* 2001; Wu and Lin 2006; Stinchcombe and Kirkpatrick 2012). Although there are multiple approaches to modeling continuous growth, one particular advantage of FVT modeling is that parameters describing developmental growth curves can be extracted from the FVT models and used as biologically interpretable and inter-relatable traits such as the relationship between growth rates, durations, inflection points, and final sizes. This ‘parameters as data’ approach enables a broad array of analyses at both genetic and phenotypic levels (Hernandez 2015; Kulbaba *et al.* 2017). In the current study, we employ a Bayesian hierarchical approach to FVT modeling that leverages global information from the entire dataset as well as each genotype to estimate replicate-level parameters describing growth curves that underlie the developmental dynamics of plant height.

One inherent but seldom addressed complication in studying developmental genetics is that development of a given trait rarely occurs independently of organism-level attributes. For instance, in plants carbon availability can severely limit and alter development, even in determinate structures such as leaves (Schneidereit *et al.* 2005; Raines and Paul 2006). Further, including physiological parameters in plant breeding models is predicted to accelerate and improve yield gains (Hammer *et al.* 2005). One solution is using a hierarchical Bayesian approach to FVT modeling that incorporates genotype-specific values for physiological conditions such as carbon availability (for instance, estimated using *A*_*max*_) to statistically factor out variation caused by resource availability. Accounting for carbon availability in FVT parameter estimation can increase estimates of heritability and improve QTL mapping results (Baker 2018a, b).

QTL mapping provides a well-tested method of uncovering the genetic architecture of Function-Valued Traits (FVT). FVT variation may arise from structural or regulatory genes that differ among sampled genotypes. Examining gene expression can therefore provide insight into the mechanistic connections between genomic architecture and developmental dynamics of phenotypes (Schmid *et al.* 2005; Li *et al.* 2010; Jiang *et al.* 2015; Zhu *et al.* 2016). We use Mutual Rank (MR) and Weighted Gene Co-expression Network Analyses (WGCNA) to identify expression networks associated with FVT trait variation. These networks are then used to focus our analysis to specific expression traits for eQTL mapping (Munkvold *et al.* 2013; Ponsuksili *et al.* 2015). Interestingly, the genomic architecture of eQTL appears to depart from that of other phenotypic QTL such as FVT QTL in two important respects: first, gene expression traits tend to have only one or a few eQTL whereas morphological phenotypic traits are often highly polygenic (Gibson and Weir 2005). Second, eQTL from multiple expression traits in diverse taxa from yeast to *Brassica* can be highly colocalized into eQTL “hotspots”. These hotspots may indicate a regulatory gene or switch that has a disproportionate impact on downstream gene expression (Schadt *et al.* 2003; West *et al.* 2007; Hammond *et al.* 2011). In contrast, QTL for morphological traits may colocalize, but typically they do not do so to the same extent (Schadt *et al.* 2003; Tian *et al.* 2016). Whether general eQTL trends hold for targeted expression traits in agroecologically relevant field settings remains unknown. Further, to the best of our knowledge eQTL mapping has not been used to examine the mechanistic basis of developmental morphology captured via function-valued trait modeling.

Here, we estimate continuous developmental growth curves of plant height, a trait that when selected upon can lead to more effective increases in yield than directly selecting on yield itself (Law *et al.* 1978), in a set of *Brassica rapa* Recombinant Inbred Lines (RILs) while mathematically factoring out the effects of carbon availability. We examine the patterns of genetic correlations among parameters describing change in height over time such as growth duration and final plant size, and we ask whether these developmental parameters correlate with yields. Using QTL mapping, we outline the genetic architecture of plant height development. Next, we use MR and WGCNA to identify genes and gene network module eigengenes whose expression patterns correlate with FVT parameters. We compare the predictive capacity of QTL and co-expression approaches in two ways: first, we test the relative effectiveness of QTL vs. MR genes vs. WGCNA module eigengenes (and combinations thereof) in explaining genetic variation of developmental traits. Second, we test whether QTL for FVT traits are enriched for genes identified via co-expression approaches. To explore the mechanistic basis of FVT QTL, we perform eQTL mapping on our MR genes and WGCNA module eigengenes. For eQTL and FVT QTL that colocalize, we explore the relative proportion *cis-* vs. *trans*-eQTL and their effect sizes. We ask whether eQTL colocalize to regulatory hotspots and if so how these compare to FVT QTL. Our eQTL analysis offers an additional line of inference for candidate gene identification as well as a potential mechanistic explanation for the regulation of yield-related FVT QTL.

## MATERIALS AND METHODS

### Species description

*Brassica rapa* (Brasssicaceae) is an herbaceous crop species first domesticated in Eurasia. This study was conducted on Recombinant Inbred Lines (RILs) derived from crossing R500, a yellow sarson oil seed variety, with IMB211, which is a rapid cycling line derived from the Wisconsin Fast Plant line (WFP). All RILs are expected to be >99% homozygous (Kokichi and Shyam 1984; Brock and Weinig 2007; Iniguez-Luy *et al.* 2009; Markelz *et al.* 2017). In comparison with IMB211, R500 flowers later, attains a larger size and greater biomass, and allocates more resources to seed production. This experiment includes 120 RILs as well as R500 and representative IMB211 genotypes.

### Experimental Design and Data Collection

In 2011, and 2012, the IMB211 × R500 RILs were germinated in the University of Wyoming greenhouse in fertilized field soil, and transplanted into the field at two planting densities, as previously described (Baker *et al.* 2015). Briefly, crowded (CR) plants consisted of 5 plants of the same genotype per 4” peat pot with the central plant designated as a focal individual. The uncrowded (UN) treatment consisted of a single plant per pot. When the cotyledons were expanded, plants were transplanted to the field into randomly located blocks that consisted of either UN or CR plants. Each block contained a full RIL set (and representatives of the RIL parental genotypes), and RIL locations were randomized within blocks with 25cm between each focal plant. For phenotypic data collection 6 UN blocks were transplanted into the field in 2011 and in 2012 8 CR and 8 UN blocks were transplanted. In 2011, an additional 5 UN blocks were transplanted into the field for RNAseq. Plants were watered daily to field capacity and treated with pesticides as needed following Baker *et al.* (2015). Each year, we collected data on the timing of germination, bolting, and flowering by surveying plants 5-7×/week. We recorded temperature data every 5s in the greenhouse and field using a series of Onset^®^ Hobo data loggers (Bourne, MA, USA) and a Campbell Scientific (Logan, UT, USA) CR23X data logger equipped with a Vaisala (Helsinki, Finland) HMP-50 sensor. Temperature data were used to produce hourly and daily means, as well as hourly and daily minimums and maximums, for Degree Day (DD) calculations, which used a *B. rapa*-specific base value of 0.96°C (Vigil *et al.* 1997).

#### Morphological data

Plant height was recorded for all plants starting at leaf emergence. In 2011, height was measured 6 times during the growing season, and these measurements captured final heights. In 2012, height was measured 2-3 times per week until senescence. Perhaps because of the increased precision in 2012 trait estimates, RNAseq data corresponds more closely to 2012 plant-level phenotypic data compared to 2011, and we focus on 2012 plant-level phenotypic data. Full analyses of FVT traits and QTL including 2011 data can be found in the supplemental materials. Flowering phenology and performance were estimated based on 2012 fruit and seed numbers, as described in Baker *et al.* (2015).

#### Function-Valued Trait (FVT) modeling and data analysis

Height data were visually inspected for erroneous data points on a replicate level following Baker et al (2015). FVT modeling for trait estimation used Bayesian approaches that fit logistic growth curves to longitudinal height data (Eqn 1; adapted from Baker *et al.* 2018a). Height for each individual replicate plant is represented by a minimum of 5 and maximum of 13 sequential measurements. Briefly, we utilized a three-level hierarchical Bayesian model that retains the measurement data structure to account for information across all plants and genetic lines within the population, including replicate plants within each line.

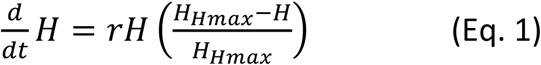

Replicate-level parameters were extracted from the fitted logistic growth curves and treated as trait data (Jaffrézic and Pletcher 2000; Kingsolver *et al.* 2001, Wu and Lin 2006; Stinchcombe *et al.* 2010; Baker *et al.* 2018a). These parameters include the growth *<u>r</u>*ate (*r*, cm/DD), and an estimate of the *<u>max</u>*imum *<u>h</u>*eight based on the asymptote of the logistic growth curve (*Hmax*, in cm). Additional parameters were algebraically extracted from the growth curve and include the *<u>d</u>*uration of growth (*d*, in DD) and the *i*nflection point of the growth curve in *<u>D</u>*egree Days (*iD*, in DD). The parameter *d* was defined as the time in DD when 95% of the final size (*Hmax*) was achieved. The parameter *iD* reflects the transition from exponentially accelerating to decelerating growth rates.

The hierarchical Bayesian model was implemented using PyMC, a Bayesian Statistical Modeling Python module. The model parameters were estimated via MCMC using the Metropolis-Hastings algorithm (Chib and Greenberg 1995; Patil *et al.* 2010). The MCMC estimations were performed using a single chain to sample 500,000 iterations, which includes the first discarded 440,000 burn-in iterations; the remaining 60,000 iterations were retained. By thinning to 1 iteration in 20, the retained iterations were reduced to 3,000 samples for every FVT parameter from which the posterior distributions were tabulated. All parameters’ trace and auto-correlation plots were examined to ensure that the MCMC chain had adequate mixing and had reached convergence. All observed data for each genotype were plotted with two 95% credible interval envelopes. The inner, yellow envelope represents the credible intervals for the model based on the observed data, and the green envelope is the 95% credible interval where future observations from the same environment are expected (Fig. 1; Kruschke 2014; Baker *et al.* 2018b).

**Fig. 1.**
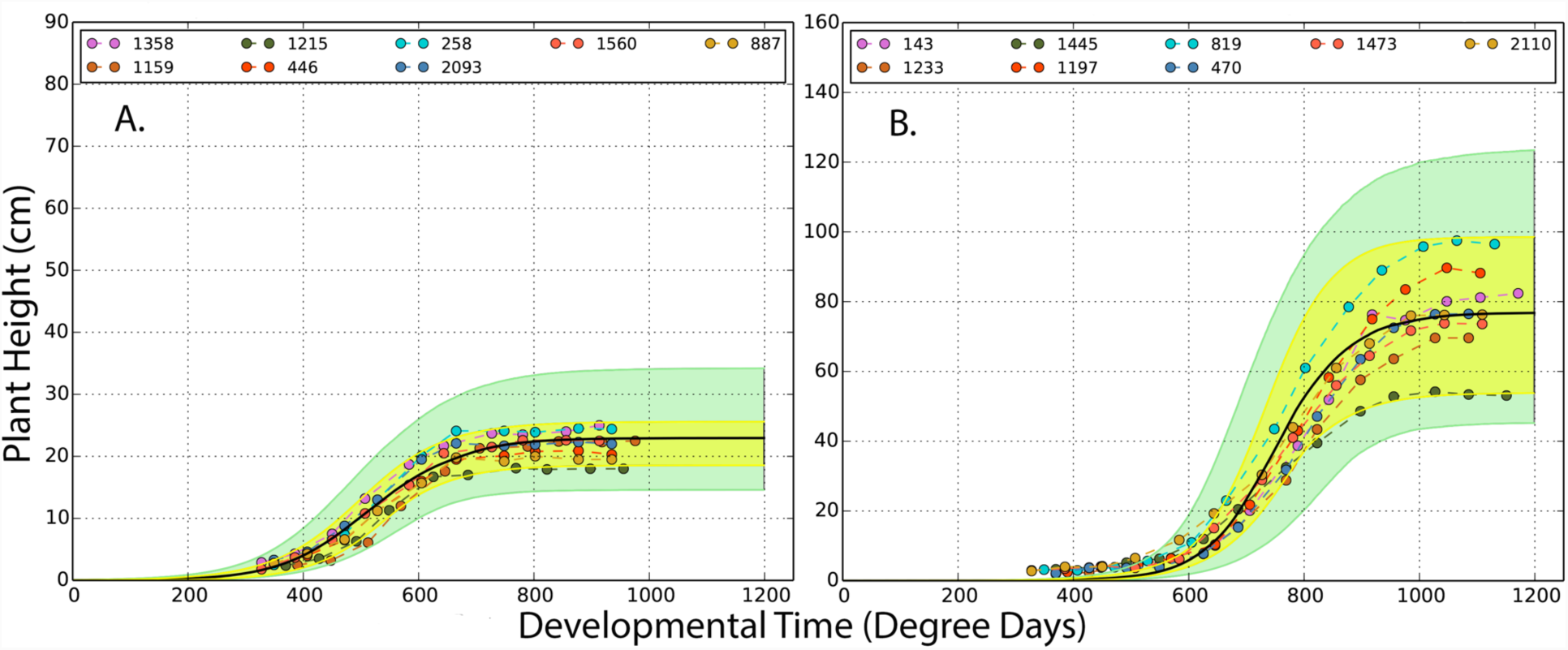
Representative genotypes (A, IMB211; B, R500) of Bayesian FVT trait estimation approaches for uncrowded plants from the 2012 season. Within each panel, dots represent observed data. Colors indicate replicates within each genotype, and indicate that each replicate was measured multiple times throughout the growing season. The black line is the Bayesian estimate of logistic growth curve that best represents each genotype. The yellow envelope is a 95% credible envelope for the observed data; the green envelope is a 95% credible envelope for where new data is predicted to occur for a specific genotype and environment combination.

#### Phenotypic plasticity

To detect environmental factors that might affect the correspondence between genotype and phenotype, we analyzed replicate level phenotypic datasets from 2012. We tested for the main effects of genotype and treatment and all possible interactions using the *lme4* and *pbkrtest* packages in the R statistical environment (Halekoh and Højsgaard 2014; R Core Team 2016; Bates *et al.* 2018). In these tests, all effects were considered random and block was nested within the treatment effect. Significant main effects of environment (treatment) were considered evidence of phenotypic plasticity, and interactions of treatment × genotype was considered evidence for genetic variation in phenotypic plasticity.

#### Best Linear Unbiased Predictions (BLUPs)

BLUPs were calculated independently for UN and CR treatments in R using the *lmer* function in the *lme4* package while controlling for block effects (Bates *et al.* 2018; Kuznetsova *et al.* 2018). Broad sense heritability (*H*^*2*^) was calculated as the genotypic variance divided by the sum of genotypic, block, and residual variances.

#### Genetic Correlations

We assessed the genetic correlations among height FVT and previously published phenology and fitness traits (Baker *et al.* 2015) across environments using Pearson’s correlations of trait BLUPs. Bonferroni corrections for multiple testing were applied to all genetic correlations.

#### QTL mapping

QTL analyses were performed in R/qtl (Broman *et al.* 2003) based on a map with 1451 SNPs having an average distance of 0.7 cM between informative markers (Markelz *et al.* 2017). The *scanone* function was used to perform interval mapping (1cM resolution with estimated genotyping errors of 0.001 using Haley Knott regression) to identify additive QTL. QTL model space was searched using an iterative process (*fitqtl, refineqtl*, and *addqtl)* to identify additional QTL while taking into account the effects of QTL identified by *scanone* and *addqtl*. All significance thresholds (0.95) were obtained using 10,000 *scanone* permutations (Broman *et al.* 2003; Broman and Sen 2009). QTL and their 1.5LOD confidence intervals are displayed using MapChart2.0 (Voorrips 2002). Percent variance explained (PVE) is calculated as PVE=100 × (1 – 10^(−2 LOD/ n)). We compared QTL peaks to the *B. rapa* genome (Version 1.5; Cheng *et al.* 2011) to identify positional candidate genes underlying each QTL. A similar approach was used for mapping eigengene QTL (see below). However, the R/qtl implementation of composite interval mapping (Broman and Sen 2009) was used.

#### RNAseq

We used the RNA sequencing data previously reported in Markelz et al (2017). Briefly, in 2011 five UN blocks of plants designated for destructive sampling were transplanted into the field and allowed to establish for three weeks. Apical meristem tissue, consisting of the upper 1cm of the bolting inflorescence, was collected from three individual replicate plants per RIL and immediately flash frozen on liquid nitrogen as described in Markelz et al (2017). RNA library preparation and sequencing were performed as previously described (Kumar *et al.* 2012; Markelz *et al.* 2017). Reads were mapped to the *B. rapa* CDS reference described in Devisetty *et al.* (2014) using BWA (Li and Durbin 2009), with an average of 6.52 Million mapped reads per replicate. Read counts were imported to R (R Core Team 2016) and filtered to retain genes where more than 2 counts per million were observed in at least 44 RILs. Libraries were normalized using the trimmed mean of M-values (TMM) method (Robinson and Oshlack 2010) and a variance stabilizing transformation was done using voom (Law *et al.* 2014).

#### Genetic network reconstruction

To reconstruct gene co-expression networks, the fitted gene expression values for each RIL from the limma-voom fit (expression ~ RIL) were used and filtered to keep the top 10,000 genes most variable between RILs.

For each sample type, two network reconstruction methods were used. First, mutual correlation rank (MR) networks (Obayashi and Kinoshita 2009) were constructed. Pairwise MRs were calculated between each of the 10,000 genes and also between each gene and the BLUP parameter estimates from the 2011 and 2012 FVT models. A series of increasingly large growth-related networks were defined using genes directly connected to the FVT parameters with MR thresholds of ≤ 10, 20, 30, and 50. Multiple different phenotypes were used to jointly seed each network, therefore networks may contain more nodes (and more genes) than the thresholds suggest. However, because some gene expression levels are uniquely correlated with specific phenotypes while others may be correlated with multiple phenotypes, the number of nodes is less than the product of the threshold value and number of phenotypes used to seed the network. Permutation analysis was used to test the network size expected by random chance at each threshold; 95 or more of 100 permutation networks had zero edges connecting FVT BLUPs and gene expression, showing that our MR networks are recovering statistically significant connections. We used the *blastn* algorithm (Altschul *et al.* 1990) with the discontiguous megablast option and an E-value cutoff of 0.001 to compare *B. rapa* genes to *Arabidopsis thaliana* genes (TAIR10 annotation; ftp://ftp.arabidopsis.org/home/tair/Sequences/blast_datasets/TAIR10_blastsets/TAIR10_cds_20101214_updated).

Second, we constructed networks using a Weighted Gene Correlation Network Analysis (WGCNA; Zhang and Horvath 2005; Langfelder and Horvath 2008). For these networks a soft threshold power of 3 was used, corresponding to the lowest power that had a correlation coefficient > 0.9 with a scale-free network topology. We used the “signed hybrid” network, which only connects genes with positive correlation coefficients. This network consisted of 50 modules with a median of 91 genes per module. The eigengene expression value of each module was determined using WGCNA functions. The Pearson correlation between each module’s eigengene expression value and each FVT BLUP was calculated to identify modules potentially related to FVTs. Modules were considered significantly associated with a FVT BLUP if the multiple-testing corrected p-value (method = “holm” in R function *p.adjust*) for the correlation test was less than 0.05. Gene Ontology (GO) category enrichment was performed on each significant module; we only examined the Biological Process (BP) and Cellular Compartment (CC) categories. Categories were considered significantly enriched if the false discovery rate adjusted p-value was < 0.05.

#### Comparing approaches for genetic architecture

We compared the effectiveness of QTL, MR, and WGCNA approaches for predicting phenotypic variation in *r* and *Hmax* through a series of multivariate linear regression models (*lm* function in R). We extracted the effect size and direction for each QTL using the *effectplot* function in r/qtl (Broman and Sen 2009). In all cases, the trait BLUPs were the dependent variable, and all allele-specific effect sizes, gene expression, and eigengene expression values were independent variables. For each trait we generated three types of additive models: 1) models with one type of independent variable (genotypic information based on alleles harbored at each QTL including allele-specific effect sizes and direction or genotype specific gene expression values for MR genes or genotype specific eigengene expression values), 2) models with two types of independent variables (QTL and MR gene expression, QTL and eigengene expression, or MR gene expression and eigengene expression), and 3) full models with all three data types as independent variables. For each trait we included only significant QTL, genes from the MR30 network, and eigengenes that were significantly correlated with the trait of interest. Each model was subjected to a backwards model reduction routine where non-significant terms were iteratively removed until all terms in the model had significant effects on the dependent variable (p<0.10). We used AIC scores to compare final models.

#### Relationships between co-expression and FVT QTL

We performed Fisher’s exact test to determine whether the FVT QTL regions were enriched for genes and/or eigengenes identified via MR and WGCNA network analyses. Enrichment of FVT QTL for MR-identified genes was interpreted as evidence that the MR-identified genes are candidate causal genes for the FVT trait of interest.

#### eQTL Analyses

To explore the regulatory mechanisms of MR-identified genes and WGCNA-identified eigengenes, as well as their potential connection to FVT QTL, we performed eQTL analyses. Our network analyses effectively allowed us to reduce the number of expression traits mapped from 10,000 to less than 75. Therefore, we used composite interval mapping (Zeng 1993), which is usually considered too computationally intensive for eQTL studies. Permutation testing (Doerge and Churchill 1996) was used to establish a p < 0.05% significance threshold for each gene. The *bayesint* function in r/qtl was used to define 99% confidence intervals for each eQTL. For some eQTL with very high LOD scores the resulting confidence interval was a single basepair (clearly unrealistic given the limitations imposed by the number of recombination events in a mapping population). For such eQTL we used a window of +/-20kb around the identified base pair as the eQTL interval. We defined *cis*-eQTL as eQTL that include the physical gene generating the mRNA transcript and *trans*-eQTL as any eQTL that does not include the physical location of the gene. For MR-identified genes, *cis*-eQTL are interpreted as evidence of variation in *cis* regulatory elements such as promoters whereas *trans*-eQTL are interpreted as evidence for *trans*-acting regulatory proteins such as transcription factors, other signaling proteins, or small RNAs that modulate gene expression. Because eigengenes represent the composite expression of a median of 90 genes, one cannot assign *cis-* vs. *trans*-eQTL identity for these traits (although the majority of their action is expected to be in trans). MR gene or eigengene eQTL that colocalize with FVT QTL may explain the underlying basis for the FVT QTL, and such colocalizing eQTL represent candidate causal genes for the FVT eQTL locus. An alternative explanation is that eQTL that co-localize with FVT QTL are in linkage disequilibrium with the FVT QTL candidate. eQTL that do not co-localize with FVT QTL may still be affecting plant development, but at a level not directly detectable in the FVT QTL mapping.

#### Data availability

The linkage map used for QTL and eQTL analyses is available in Markelz et al (2017). Replicate level FVT parameters are presented in S1; RIL-specific gene expression values will be made available in supplemental materials (via FigShare) upon acceptance of the manuscript and are available to the editor and reviewers upon request.

## RESULTS

### FVT Modeling

For all FVT modeling, the data were sufficient to support all aspects of the growth curves modeled, and the models fit the data well (Fig. 1 for example model fits). Plots for all FVT models can be found in S2.

### Phenotypic Plasticity and Heritability

To assess the effects of the environment on plastic growth responses, we analyzed raw replicate level data. Although there were main effects of Block (nested within treatment) and genotype (RIL ID) for all traits, there were no significant main effects treatment (Table 1). However, there was genetic variation for a plastic response to crowding for all traits except *iD* (inflection time, in <u>D</u>egree Days; treatment-by-genotype interaction; Table 1).

**Table 1.** Phenotypic plasticity and heritabilities of FVT parameters. Block is nested within the Treatment effect. Treat corresponds to the crowded and uncrowded treatments in 2012 and Genotype indicates RIL id. Significant effects are emphasized by bold text.

| Trait | Model t-value (df) | Random effects – Chi Square value (degrees of freedom) |  |  |  | Heritabilities (%) |  |
| --- | --- | --- | --- | --- | --- | --- | --- |
|  |  | Block (Treat) | Treat | Geno-type | Treat* Geno-type | UN 2012 | CR 2012 |
| ***R*** | **16.62 (1.08)**<br>* | **80.2 (2)**<br>*** | 7.28e-12 (1)<br>NS | **136 (1)**<br>*** | **211 (1)**<br>*** | 74.5 | 76.0 |
| ***D*** | **43.32 (1.57)**<br>** | **58.5 (2)**<br>*** | 3.64e-12 (1)<br>NS | **294 (1)**<br>*** | **4.88 (1)**<br>* | 79.5 | 79.3 |
| ***iD*** | **37.16 (1.65)**<br>** | **98.2 (2)**<br>*** | 1.42e-10 (1)<br>NS | **369 (1)**<br>*** | 0.34 (1)<br>NS | 86.8 | 83.7 |
| ***Hmax*** | **8.70 (1.83)**<br>* | **116 (2)**<br>*** | 0.0 (1)<br>NS | **226.4 (1)**<br>*** | **42.3 (1)**<br>*** | 81.2 | 68.1 |

Signif. codes: p < 0.001 '\*\*\*'; p < 0.01 '\*\*'; p < 0.05 '\*'; p < 0.1 '.'; p > 0.1 'NS'

In general, heritabilities were higher for plants grown in the UN relative to CR treatments for all traits. This may reflect the relatively stochastic nature of the crowding response: in some cases in the CR treatment the focal plant may have outcompeted its neighbors whereas in others it may have been outcompeted.

### Genetic Correlations

To explore the genetic relationships among the height FVT parameters and previously published estimates of plant phenology and fitness, we conducted a correlation analysis on BLUPs of each trait. In general, the pattern of genetic correlations within years and treatments was similar. UN*r* from 2012 was correlated with all traits except *Hmax* (Fig 2). In contrast, CR*r* in 2012 was negatively correlated with other all other 2012 CR FVT traits, with all CR phenology traits (except the bolting-to-flowering interval) and CR fitness traits (S3). UN*r* in 2012 was negatively correlated with UN*d* and *iD* but not *Hmax*. UN*r* 2012 was also negatively correlated with phenology and fitness. These patterns of genetic correlations are largely consistent across years and treatments; a representative subset are presented in Fig 2.

**Fig. 2.**
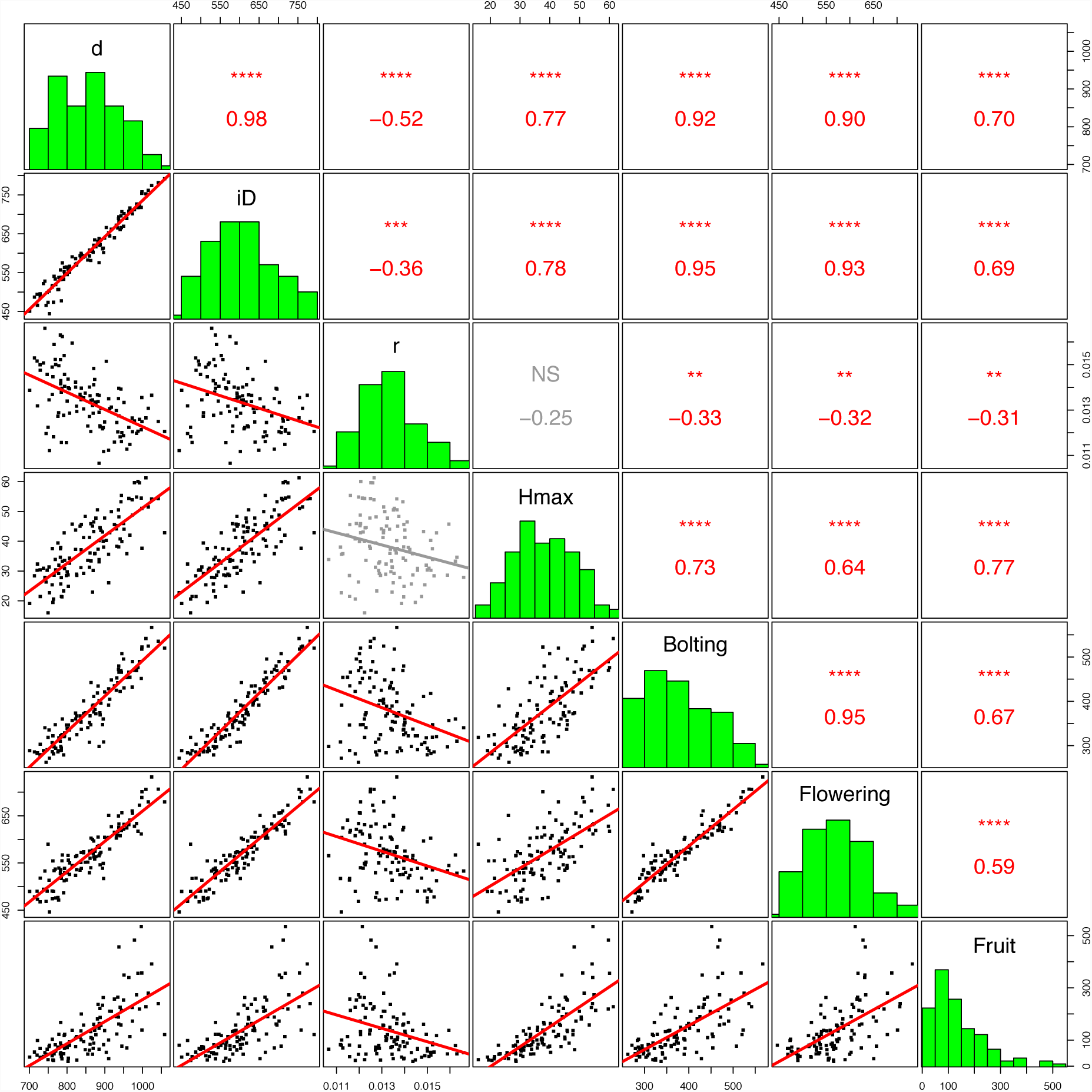
Genetic correlations among UN 2012 FVT height, phenology, and fitness traits. Each point is a genotypic mean (BLUP). Bonferroni corrections for multiple tests (n=7) have been applied. Non-significant correlations are in gray. All time is expressed in Degree Days. * p<0.05, ** p<0.01, *** p<0.001, **** p<0.0001, NS p ≥ 0.05.

### QTL mapping

To further explore the genetic architecture of the height FVT parameters, we conducted QTL mapping analyses of the height FVT traits. In total we mapped 32 individual QTL from 2012 (2011 FVT QTL are presented in S4); however, an alternative interpretation is that we mapped as few as 9 highly pleiotropic QTL. QTL were observed throughout the genome, except on chromosomes 2, 4, and 8. Most QTL localized to chromosome 3, 9 and 10. Across all traits, each QTL explained 29% of trait variation on average. The minimum explained variance was 9.5% and the maximum was 73% of variance (Fig 3 & S4).

**Fig. 3.**
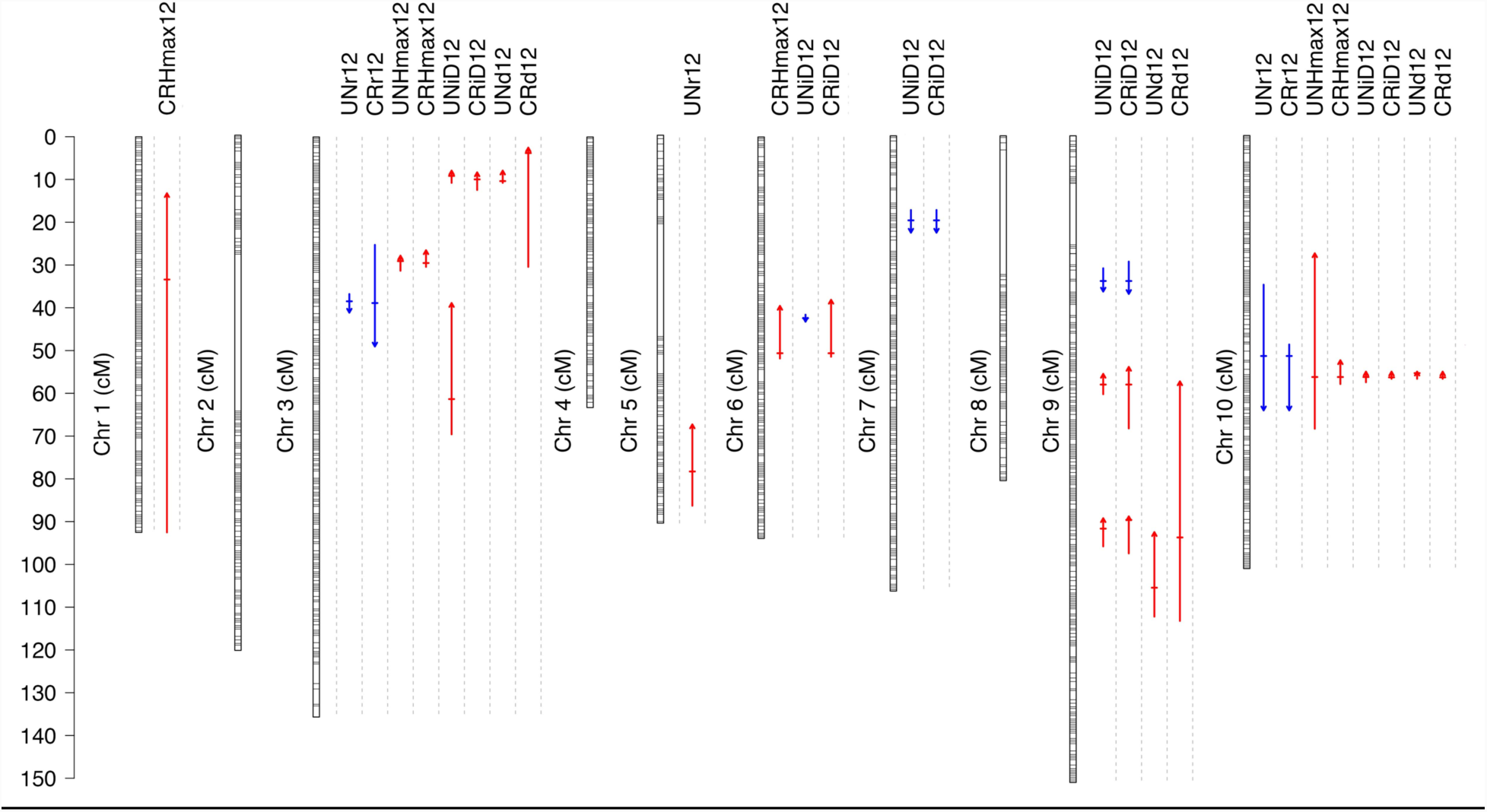
A map of all QTL identified in 2012. Horizontal lines on chromosomes indicate the position of RNAseq markers used to genetic map construction. Each QTL is indicated with a vertical arrow under the trait name. Horizontal hatches indicate QTL position, the arrow length indicates 1.5 LOD support limits. Arrow heads and color (up, red = positive; down, blue = negative) indicate QTL direction relative to the R500 parent. Exact locations, markers, and LOD scores for all QTL can be found in S4.

### Genes under FVT QTL

To determine positional candidates within mapped FVT QTL, we compared our FVT QTL to the *B. rapa* genome and identified genes underlying the QTL. We restricted our search to QTL with LOD > 9 (Table 2). All 9 of these QTL were on either chromosome 3 or 10. Because several of the QTL co-localized (had overlapping 1.5 LOD confidence intervals), we often found the same genes under multiple QTL. After removing duplicate entries, we found 490 unique genes underlying the 9 QTL investigated (S5).

### RNAseq

We used RNA sequencing (RNAseq) to understand the genetic mechanisms underlying FVT QTL and as an alternative approach for examining the genetic architecture of our FVT traits without *a priori* knowledge. 21,147 genes of 28,668 genes with detectable expression in UN treatment were differentially expressed among RILs (FDR < 0.01). The 10,000 genes with the most variable expression among RILs were used for downstream network analysis.

### Mutual Rank Network Analysis

To find gene co-expression networks relevant to the FVT model parameters, we built Mutual Rank (MR) networks nucleated on each FVT model parameter and performed permutation analyses to determine the statistical significance of our networks. Ninety-five or more of 100 permutations had zero connections between FVT parameters and gene expression. Therefore, our MR networks are enriched for *bona fide* connections at a variety of MR threshold cutoffs (The MR30 network is shown in Fig 4; larger networks become difficult to visualize and are presented in S6). Complete gene membership for all MR-thresholds annotated with the best hit obtained by *blastn* against the predicted *A. thaliana* proteome are presented in supplemental materials S7.

**Fig. 4:**
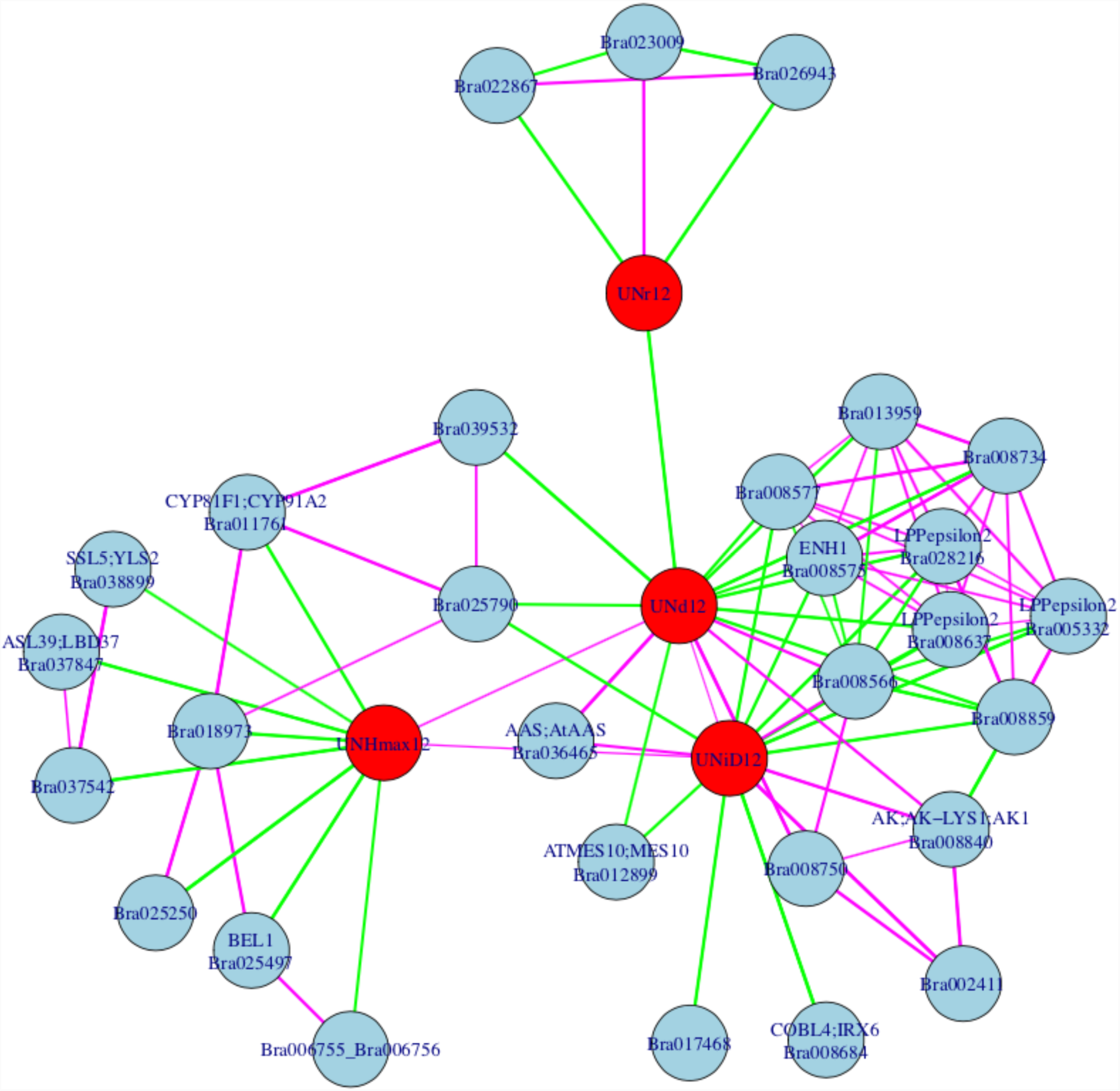
A scale-free diagram of the Mutual Rank network nucleated around FVT traits from 2012 with a cutoff of 30. Network nodes consist of either FVT traits or co-expressed genes. FVT traits are shown in red circles and genes are indicated in blue circles. Network edges indicate significant correlations. Purple lines indicate positive correlation values while green lines indicate negative correlation values and line thickness corresponds to strength of the correlation. UN, uncrowded; *r*, growth rate; *d*, duration of growth; *iD*, time in degree days when the growth curve reached its inflection point; *Hmax*, estimated maximum height based on FVT modeling. Additional network cutoffs, 2011, and 2012 crowded networks are in S6; gene names and annotations are in S7.

We used Fisher’s exact test to determine whether FVT QTL were enriched for MR-identified genes. We found no evidence for enrichment for MR10 networks (p=1.0) but significant evidence for enrichment for MR20, MR30, and MR50 networks (p<5E-09; Table 4). In theory, MR10 networks should contain only those genes whose expression values are most highly correlated with FVT phenotypes. The non-significant results for MR10 may be caused by low power due to the single gene identified.

**Table 4.** Fishers exact tests for enrichment of FVT QTL for MR-identified genes.

|  |  | QTL |  | p-value |
| --- | --- | --- | --- | --- |
|  |  | Yes | No |  |
| **MR10** | Yes | 0 | 2 | 1.0 |
|  | No | 5,816 | 37,645 | (NS) |
| **MR20** | Yes | 16 | 0 | 6.91e-13 |
|  | No | 6,800 | 37,647 | *** |
| **MR30** | Yes | 25 | 4 | 4.98e-09 |
|  | No | 5,791 | 37,643 | *** |
| **MR50** | Yes | 46 | 10 | 9.93e-21 |
|  | No | 5,770 | 37,637 | *** |

p > 0.05, NS; p < 0.0001, \*\*\*\*

### Weighted Gene Co-expression Network Analysis (WGCNA)

In a second approach to identifying gene expression networks related to estimates of FVT trait parameters, we used a Weighted Gene Co-expression Network Analysis (WGCNA) to identify eigengene modules. Modules of interest were identified as those showing a significant correlation between eigengene expression values and FVT model parameters across the RILs (Figure 5). Gene Ontology (GO) enrichment analysis was performed to examine the potential function of correlated module (S8); below we discuss correlations with modules that had at least one GO term enriched. There are positive correlations between 2012 BLUPs for maximum height (*Hmax*), growth *duration* (*d*), and the time that the growth curve reached its inflection point (*iD*) and the *cyan* module (related to protein translation), the *midnight blue* module (related to wounding/herbivore defense responses as well as some abiotic stress responses), and the *blue* module (enriched for genes related to cell division and development). This suggests that plants that have a longer duration of growth and reach a higher maximum height are producing more protein, undergoing more rounds of cell division, and have increased defense signaling. These three parameters also showed negative correlation with the brown module (enriched for actin cytoskeleton and protein dephosphorylation terms). *Hmax* is negatively correlated with *yellow* (enriched for terms related to photosynthesis). This correlation could be caused by a difference in cellular maturation rates: plants with more rapid cellular differentiation would be expected to show an upregulation of chloroplast genes and reduced growth due to earlier differentiation and consequently relative lack of cell elongation.

**Fig. 5.**
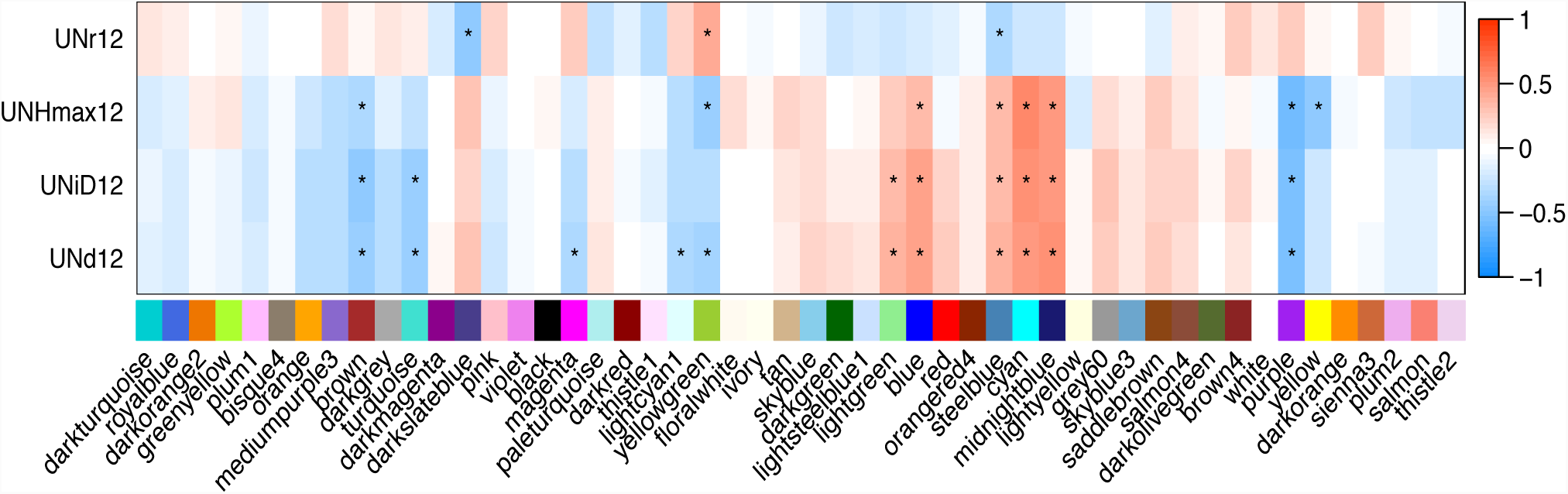
Correlations among WGCNA identified eigengenes and UN 2012 FVT traits. Significant correlations are denoted with an asterisk. *r*, growth rate; *d*, duration of growth; *iD*, time in degree days when the growth curve reached its inflection point; *Hmax*, estimated maximum height based on FVT modeling.

#### Comparisons of QTL and network modeling for phenotypic prediction

To compare the effectiveness of various approaches and combinations of these approaches in explaining the variation in FVT trait estimates, we compared a series of additive linear models based on QTL, MR genes, or WGCNA eigengenes both singly and in combination. For UN*r* (in 2012), models containing only QTL outperformed models containing either MR30 identified gene expression or WGCNA-identified eigengene expression (Table 5). For two-data type models, models with only QTL outperformed those containing multiple data types. For *Hmax*, MR gene expression outperformed both QTL and WGCNA-identified eigengene expression as well as combinations of two data types. For both traits, the full model (with all three data types for *r*, but which reduced to WGCNA and MR gene expression values for *Hmax*) were the best models for explaining phenotypic variation (*r*: F_(5,110)_=25.31, p<0.0001; *Hmax*: F_(9,106)_=33.16, p<0.0001). Similarly, the best two-data type models were a significantly better fit to the data than the best single-data type models (*r*: F_(5,114)_=40.182, p<0.0001; *Hmax*: F(4,113)=80.398, p<0.0001). For all comparisons, the significantly better model according to ANOVA also had lower AIC scores (Table 5). Taken together, these results indicate that although each approach has significant predictive capacity, combining multiple approaches improves estimation of trait variation.

**Table 5.** Comparison of additive linear models using genetic and transcriptomic data to explain 2012 uncrowded phenotypic data.

| Trait | Best single-data type model | AIC | Next best AIC (next best model) | Formula<sup>§</sup> | Best model F-value (DF), significance and adjusted R<sup>2</sup> |
| --- | --- | --- | --- | --- | --- |
| *r* | QTL | -1305.97 | -1256.43 (WGCNA) | $y \sim rQTL2 + rQTL2 + rQTL3$ | F(3, 113)= 30.9 ***<br>R<sup>2</sup>=0.4361 |
| *Hmax* | MR | 735.5348 | 783.6546 (WGCNA) | $y \sim Bra03899 + Bra011761 + Bra006755\_Bra006756 + Bra036465 + Bra008859 + Bra037542$ | F(6,109)=45.48 ***<br>R<sup>2</sup>=0.6989 |
| **Best 2-data type model** |  |  |  |  |  |
| *r* | QTL + WGCNA (reduces to just QTL) | -1305.97 | --1297.869 (MR+WGCNA) | $y \sim rQTL1 + rQTL2 + rQTL3$ | F(3,113)=30.9 ***<br>R<sup>2</sup>=0.4361 |
| *Hmax* | MR + WGCNA (reduces to just MR) | 734.2895 | 752.3889 (QTL+MR; reduces to just MR*)<sup>†</sup> | $y \sim Bra011761 + Bra006755\_Bra006756 + Bra13959 + Bra08840 + Bra008859 + Bra037542\_Bra002411$ | F(7,108)=40.16 ***<br>R<sup>2</sup>=0.7045 |
| **Best overall model** |  |  |  |  |  |
| *r* | Full model (QTL+ MR+ WGCNA) | -<br>1308.602 | -1305.97 (QTL + WGCNA) | $y \sim rQTL2 + yellowgreen + Bra006755\_Bra06756 + Bra025790 + Bra028216$ | F(5, 110)=25.31 ***<br>R<sup>2</sup>=0.5138 |
| *Hmax* | Full model (reduces to MR + WGCNA) | 731.63 | -734.2895 (MR + WGCMA; reduces to just MR*) | $y \sim yellow + Bra011761 + Bra006755\_Bra006756 + Bra008575 + Bra008577 + Bra008840 + Bra008859 + Bra037542 + Bra002411$ | F(9,106)=33.16 ***<br>R<sup>2</sup>=0.7157 |

\*\*\* p < 0.0001
\* This model reduced to include just MR gene expression values but is different from the best Hmax single-data type model that also includes just MR gene expression values.
<sup>§</sup> rQTL 1-3 have markers at A03x 6417941, A05x23393567, and A10x11427369, respectively,

### eQTL analyses and colocalization of eQTL with of FVT QTL

Because including MR and WGCNA results both improved upon linear models for FVT traits that contained just QTL (Table 5) and because all models that included MR and WGCNA gene/eigengene expression values were significant and predicted FVT trait variation, we used eQTL analyses to assess the mechanistic relationship between gene/eigengene expression and FVT QTL. For the 29 MR30-identified genes, we found significant eQTL on all chromosomes except 5 and 8. In congruence with FVT QTL mapping results, there were eQTL with particularly high LOD scores on chromosomes 3 and 10 (LOD >75; Figure 6). There was significant overlap among 2012 FVT-QTL confidence intervals and MR-eQTL confidence intervals based on permutation tests (n=1000, p=0.003).

**Fig. 6.**
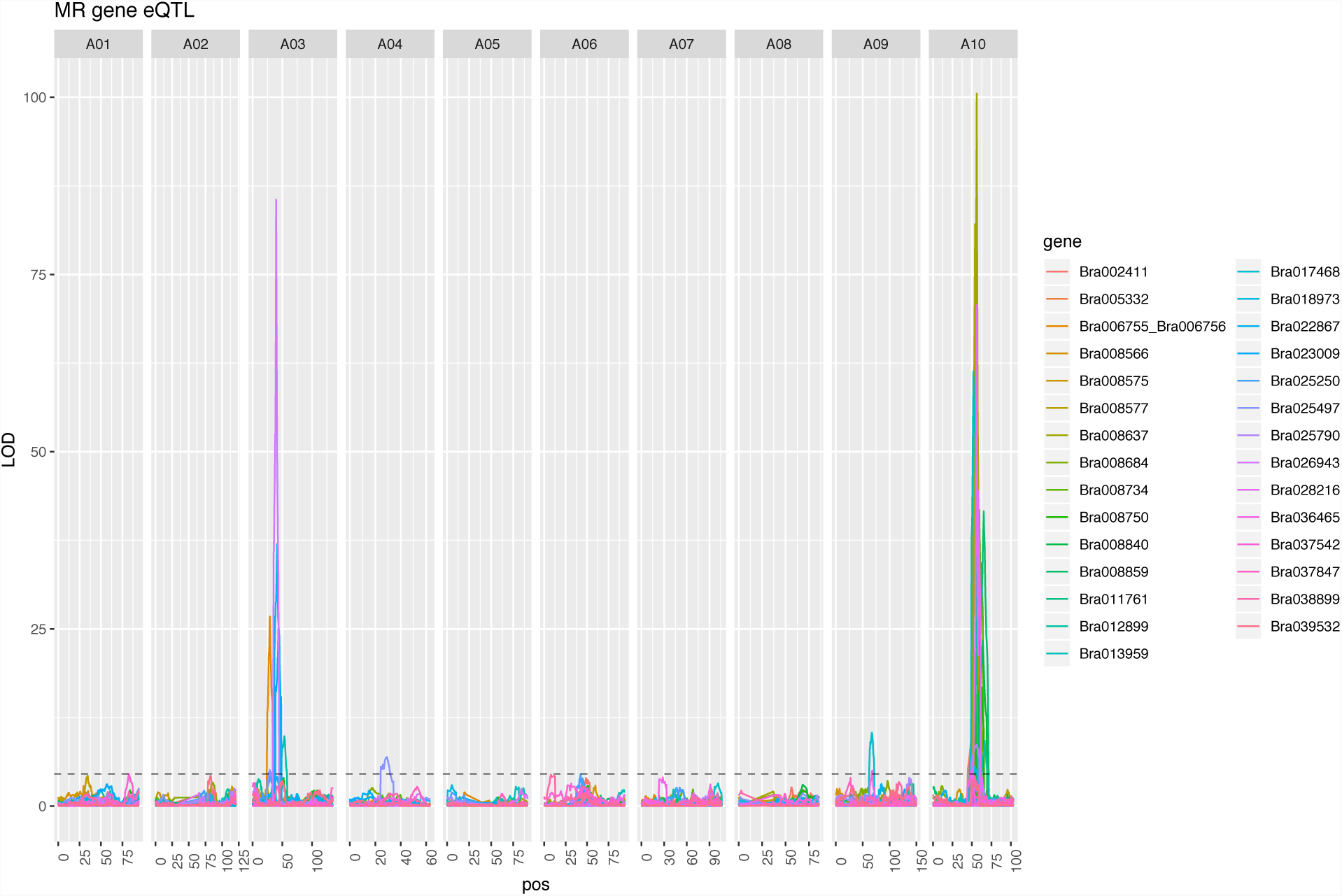
Expression trait QTL (eQTL) identified using Composite Interval Mapping (CIM) for MR30-identified genes where MR networks were nucleated around UN FVT traits. Note the eQTL hotspots on chromosomes 3 and 10.

Of the 57 MR50 genes, 42 genes had a total of 47 eQTL that overlapped with FVT QTL with LOD scores ranging from 100.5-4.6. Six of the 42 MR50 genes with eQTL that colocalized with FVT QTL had *cis*-eQTL, and of those six, three were in networks with cutoffs of MR30 or below (Table 6). The co-occurrence of these loci as MR-identified *cis*-eQTL and FVT QTL indicates that they are strong candidate genes for regulating the FVT traits. For any given FVT trait, none of the MR genes with *cis*-eQTL also had *trans*-eQTL that colocalized with other FVT QTL. Of the 36 MR genes with *trans*-eQTL that colocalized with FVT QTL, 33 had a single *trans*-eQTL that colocalized with FVT QTL. Three genes (Bra012899, Bra014655, and Bra029573) had *trans*-eQTL that colocalized with two or more distinct FVT QTL (Table 7).

**Table 6.** MR-identified genes with *cis*-eQTL that co-localize with UN 2012 FVT QTL. Note that because FVT QTL overlap a single MR *cis*-eQTL may colocalize with FVT QTL for multiple traits.

| MR gene | MR Network | Chromo-some | FVT trait | eQTL LOD range | AGI | *A. thaliana* symbol |
| --- | --- | --- | --- | --- | --- | --- |
| Bra008840 | 20 | 10 | *r, Hmax* | 22.526-23.419 | AT5G13280 | AK;AK-LYS1;AK1 |
| Bra008859 | 20 | 10 | *r, Hmax* | 41.593-41.593 | AT5G13070 | NA |
| Bra008750 | 30 | 10 | *r, iD, Hmax* | 15.375-17.671 | AT5G14600 | NA |
| Bra008711 | 50 | 10 | *r, Hmax* | 24.585-26.594 | AT5G15250 | ATFTSH6;FTSH6 |
| Bra008931 | 50 | 10 | *r, Hmax* | 8.515-10.223 | AT5G11880 | NA |
| Bra029100 | 50 | 3 | *r* | 28.288-30.055 | AT5G53045 | NA |

**Table 7.** MR-identified genes with multiple *trans*-eQTL that co-localize with FVT QTL.

| MR gene | MR Network | Chromosome | FVT Trait | eQTL LOD range |
| --- | --- | --- | --- | --- |
| Bra012899 | 10 | 3 | *iD* | 8.315-9.854 |
|  |  | 10 | *Hmax* | 6.970-9.214 |
| Bra014655 | 50 | 3 | *r, iD, d, Hmax* | 2.857-4.930 |
|  |  | 6 | *iD* | 6.526-8.333 |
|  |  | 10 | *r, Hmax* | 3.290-5.176 |
| Bra029573 | 50 | 3 | *Hmax* | 5.096-7.279 |
|  |  | 6 | *iD* | 3.052-5.367 |
|  |  | 10 | *Hmax* | 2.906-4.812 |

Next we performed eQTL analyses (Figure 7) for the 11 WGCNA-identified eigengene modules based on UN 2012 FVT (see Figure 5). Chromosome 3 harbored strong eQTL for “darkslateblue”, “steelblue”, and “yellowgreen” (all with no go enrichment; nge). Chromosome 6 had QTL for “blue” (cell division), “cyan” (translation), and “midnightblue” (herbivore/wounding). Chromosome 10 had Eigengene eQTL in two locations, one for “brown” (actin cytoskeleton) and “lightgreen” (nge), the other for “cyan” (translation), “midnightblue” (herbivore/wounding), “turquoise” (nge), and a suggestive peak for “blue” (cell division). Five of the eleven eigengenes had eQTL also colocalized with FVT QTL, indicating a potential causative connection between eigengenes and FVT for *r, iD*, and *Hmax* (Table 8). However, each eigengene had only one eQTL that colocalized with an FVT QTL.

**Table 8.** Eigengene eQTL and FVT QTL colocalization.

| Trait (eigengene) | Chromosome | FVT trait | LOD range |
| --- | --- | --- | --- |
| brown | 10 | *r*, *Hmax* | 6.052-7.223 |
| cyan | 10 | $iD, r, Hmax$ | 4.900-5.880 |
| darkslateblue | 3 | $r, iD$ | 47.828-47.828 |
| midnightblue | 10 | $r, Hmax,$ | 8.826-9.674 |
| turquoise | 10 | $r, Hmax$ | 3.967-5.840 |
| yellowgreen | 3 | $Hmax$ | 48.424-48.550 |

**Fig. 7.**
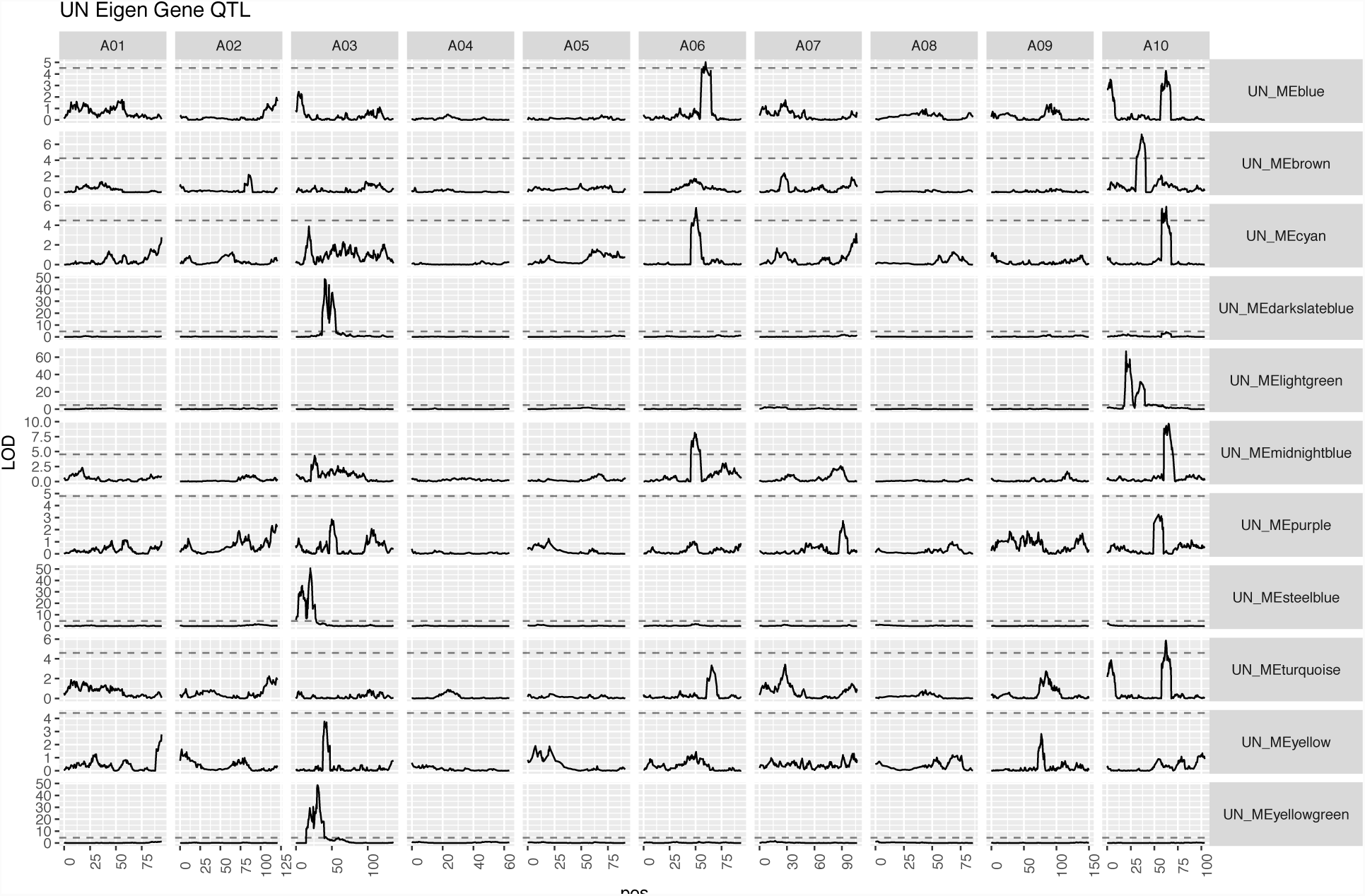
Expression trait QTL analysis (eQTL) for WGCNA-identified eigengenes that significantly correlate with UN FVT traits.

The second chromosome 10 location (“cyan”, “midnightblue”, and “turquoise”) overlaps with the FVT QTL9 and the Eigengenes has significant correlations with *d* and *iD* FVTs indicating a possible causative connection. We then performed permutation tests and determined that FVT-QTL were enriched for WGCNA-eQTL (n=1000, p=0.005).

## DISCUSSION

Plant height is often correlated with fitness and yield. Height is a complex and dynamic trait that changes over the course of development, and variation in plant height is necessarily generated through variation in developmental dynamics. However, similar heights can be achieved through multiple different growth curves. Quantifying the underlying genetic architecture and mechanistic basis of growth dynamics may result in improved estimations of final plant height, fitness, and yield. Here, we use Bayesian hierarchical modeling to estimate Function-Valued Trait (FVT) parameters describing continuous plant growth and explore their correlations with phenology and fitness. We test whether QTL mapping, genes identified through Mutual Rank (MR) co-expression, or eigengenes identified through Weighted Gene Network Co-expression Analysis (WGCNA) co-expression, or combining these information types best explain genetic variation in agroecologically relevant FVT traits in the field. Further, we employ eQTL analyses to explore the molecular genetic regulatory mechanisms that mechanistically connect FVT QTL with phenotypic variation.

Although development typically occurs in a continuous fashion, most studies quantifying development necessarily collect data at discrete timepoints. We take a “parameters as data” approach to FVT modeling to estimate the continuous nature of plant development (Hernandez 2015; Kulbaba *et al.* 2017). Much as floral development or leaf development has well defined core molecular genetic pathways that govern organ formation, elaboration, or elongation (reviewed in Bowman *et al.* 2012), there is likely a core genetic architecture that contributes to plant height. However, exogenous and endogenous factors can influence the outputs of these developmental programs. For instance, crowding may trigger a shade avoidance response and lead to rapid increases in height (e.g. Schmitt *et al.* 2003). Similarly, plant carbon status can affect the developmental morphology and final size of organs such as leaves (Schneidereit *et al.* 2005; Raines and Paul 2006; Baker *et al.* 2018a). We took two approaches to examining the core developmental genetics of plant height. First, we grew plants across multiple growing seasons and in crowded and uncrowded conditions. Second, we included a genotype-specific co-factor in our FVT models that accounts for variation in photosynthetic rates (approximated through *Amax*), thereby statistically factoring out variation due to carbon availability and allowing us to more directly interrogate the developmental genetic architecture and molecular mechanisms contributing to plant height (Baker *et al.* 2018a; b). In our study, all FVT traits had relatively high broad sense heritabilities (>70%), and all had significant main effects of genotype. Interestingly, although there were no significant main effects of treatment (i.e. population means did not differ), all FVT trait estimates (except *iD*) exhibited genetic variation for carbon-independent phenotypic plasticity via a treatment-by-genotype interaction, likely because of rank-order differences across treatments at the genotypic level (Table 1).

Morphological phenotypes, such as components of yield and height, can be highly integrated throughout development (reviewed in Klingenberg 2014). Final height is often used as a proxy for yield or fitness, yet plant growth dynamics throughout ontogeny may also be correlated with aspects of yield such as fruit and seed set (Yin *et al.* 2011; Tanger *et al.* 2017). In our experimental set of *Brassica rapa* Recombinant Inbred Lines (RILs), plant developmental dynamics including duration of growth (*d*), the inflection point in the growth curve that represents the change from exponentially accelerating to decelerating growth (*iD*), and estimates of final plant height (*Hmax*) were all significantly and positively genetically correlated (Fig 2). Interestingly, growth rates (*r*) were negatively correlated with *d* and *iD*, but were not correlated with *Hmax*, indicating that while there is a trade-off between growth rates and durations, duration of growth may be more important for final plant height than growth rate. All of our estimates of plant growth and final size were significantly genetically correlated with both phenology and yield traits. The significant correlations of *r* with yields indicates that developmental dynamics of a given trait can be related to crop yields and plant fitness through mechanisms that may be at least partially independent of final size. Because final size is positively correlated with yields while growth rates are negatively correlated with yields, selection for maximum yields at early harvest dates may come at the expense of late harvest yields and vice versa.

To examine the genetic architecture underlying the FVT estimates of growth rates, durations, and final sizes, we used standard QTL mapping procedures, which revealed a number of QTL. Of particular note, when QTL for *r* colocalized with *d*, the QTL were of opposite sign, confirming our negative genetic correlations between growth rates and durations, and indicating potentially pleiotropic loci contributing to both traits. On average, FVT QTL explained 24% of trait variation and the number of genes under each QTL ranged in to the hundreds. In part to narrow down the list of candidate genes and in part to understand the mechanistic regulation of FVT via QTL, we took two additional transcriptomic co-expression approaches to exploring the genetic architecture of FVT traits: First, we seeded a Mutual Rank (MR) co-expression network with FVT traits and asked which gene expression values correlated with variation in FVT traits. Second, we constructed 50 eigengenes based on a Weighted Gene Co-expression Network Analysis (WGCNA) and asked which eigengenes were correlated with individual FVT trait. We found that FVT QTL were significantly enriched for MR genes, indicating that these two approaches were identifying some common drivers of FVT traits. To compare the effectiveness of all three approaches, we asked whether QTL, MR genes, or eigengenes best explained variance in FVT traits. Although QTL outperformed both co-expression network modeling approaches for *r*, combining data from multiple approaches yielded improvements in our models, indicating that even though QTL, MR genes, and eigengenes often physically co-localize within the genome, they are not synonymous with one another (Table 5).

To better understand the potential function of genes related to growth WGCNA and MR networks, we used gene annotations and homology to *A. thaliana*. Although about half of the eigengenes that correlated with FVT BLUPs had no gene ontology enrichment, three eigengenes with eQTL on chromosome 10 were enriched for actin/cytoskeleton, herbivore/wounding and cell division, respectively. The MR30 genes include a homolog of the homeodomain gene *BEL1* (*NACA3* (Reiser *et al.* 1995) which is negatively correlated with *Hmax*); *BEL1* homologs have been implicated in regulation of the shoot apical meristem (Rutjens *et al.* 2009) and thus could be related to plant growth. An additional gene was identified with homology to the COBRA family gene *COBL4/IRX6* (negatively correlated with *iD*), involved in secondary cell wall biosynthesis. The MR30 network also contains a number of genes involved in metabolic homeostasis. Four of these genes are localized to the plastid and negatively correlated with *d* and *iD*, including three orthologs of the *plastidic lipid phosphate phosphatase epsilon 2* gene (*LPPε2*), which is potentially involved in synthesis of diacylglycerol, a precursor to essential photosynthetic membrane components (Nakamura *et al.* 2007). Another plastid-localized MR30 network gene is *ENHANCER OF SOS3-1 (ENH1)*; *ENH1* functions to mitigate the effects of reactive oxygen species (Zhu *et al.* 2007). Thus, plants with longer growing periods appear to put less resources into photosynthesis. The MR30 network also includes a homolog of the *A. thaliana LATERAL ORGAN BOUNDARY DOMAIN37 (LBD37)* gene, an important regulator of nitrogen response in both *A. thaliana* and *Oryza sativa* (Rubin *et al.* 2009; Albinsky *et al.* 2010). *LDB37* is negatively correlated with *Hmax.* Two genes involved in amino acid synthesis or homeostasis are present in the MR30 network and show positive correlations with *d* and *iD*: a homolog of *ASPARTATE KINASE1 (AK1),* required for regulation of aspartate, lysine, and methionine (Clark and Lu 2015), and *AROMATIC ALDEHYDE SYNTHASE (AAS)*, which converts phenylalanine into phenylacetaldehyde (Gutensohn *et al.* 2011). Overall the MR30 network results point to a close connection between metabolic regulation and growth.

Transcriptomic data allowed us to further explore the regulatory control of the FVT using eQTL mapping of WGCNA eigengenes and MR genes. eQTL mapping treats gene expression levels as quantitative traits. When combined with QTL studies of morphological phenotypes, the ultimate goal of eQTL mapping is to identify the molecular genetic changes in gene expression that lead to structural phenotypic variation, thus providing mechanistic explanations for the associations between genotype and phenotype (Schadt *et al.* 2008). In humans, such studies demonstrate that eQTL can be used in a cell-type specific fashion to annotate GWAS associations (Brown *et al.* 2013). In our study, 42 MR genes had eQTL that colocalized with FVT QTL and 6 of the 11 WGCNA eigengenes that correlated with FVT also had eQTL that colocalized with FVT QTL. These data demonstrate that the relationship between genomic loci (FVT QTL) and phenotypic variation in FVT traits is likely mediated by gene expression, specifically the expression of the genes and eigengenes we identified via MR and WGCNA.

Our eQTL results qualitatively departed from common morphological trait QTL analyses in two ways. First, MR-identified gene expression traits mapped to all chromosomes except chromosome 2, but two locations had multiple eQTL with very high LOD scores (>75): the top of chromosome 3 and the middle of chromosome 10. Virtually all genes had eQTL that mapped to one of these two locations, a common result potentially indicating an eQTL ‘hotspot’. A previous study of the effects of soil phosphorous using the same *B. rapa* RILs also identified eQTL hotspots (Hammond *et al.* 2011), but on different chromosomes. The colocalization of eQTL hotspots and FVT QTL may indicate novel regions involved in pleiotropic co-regulation of several downstream genes in the regulatory network contributing to change in plant height (Gibson and Weir 2005).

Although the presence of eQTL hotspots indicates pleiotropic gene regulation, our eQTL analyses also qualitatively departed from the FVT QTL analysis in that most of the gene expression traits we mapped were not polygenic. Of the 42 MR gene expression traits mapped, only three had eQTL that colocalized with more than one FVT QTL. eQTL studies commonly find a relative paucity of polygenic regulation compared to structural QTL studies, and our results support the general consensus that expression traits and structural phenotypes have distinctly different genetic architectures (but see West *et al.* 2007 for a counter-example). However, most eQTL are of relatively large effect, meaning that many small effect eQTL could remain undetected and contribute to polygenic regulation of gene expression traits (Gibson and Weir 2005), and these eQTL may or may not occur in regulatory hotspots.

To further understand the regulation of expression traits and FVT QTL, we divided MR eQTL into two classes: putative *cis-* and *trans*-eQTL where *cis*-eQTL likely correspond to *cis*-regulatory elements influencing gene expression (Doss *et al.* 2005). In contrast, *trans*-eQTL do not contain the gene whose expression pattern is mapped and likely correspond to *trans*-acting factors such as transcription factors that influence the MR gene expression (Hansen *et al.* 2008). In our study, of the 42 MR genes with eQTL that colocalized with FVT QTL, only five were in *cis* and the remaining 37 were in *trans*, which is only slightly higher than the proportion of *trans-*eQTL identified in an intraspecific maize cross (Swanson-Wagner *et al.* 2009). Because the *rapa* RILs are also generated from an intraspecific cross, our results are consistent with theoretical and experimental work suggesting that *trans* gene regulation should be more prevalent than *cis* regulation at the intraspecific level (Wittkopp *et al.* 2008; Goncalves *et al.* 2012, but see O’Quin *et al.* 2012 for an exception). Although unlikely given the genetic architecture of our eQTL, biases towards *trans* regulation may also stem from highly pleiotropic genes (reviewed in Signor and Nuzhdin 2018). Other authors have offered an alternative interpretation: in *A. thaliana* the proportion of *cis*-to *trans*-eQTL appears to scale with statistical power and the ability to detect small effect eQTL. *Tran*s-eQTL are typically assumed to be of small effect and so increasing sample size, replicate number, or density of markers on the genetic map should in theory increase the proportion of *trans*-eQTL detected (Hansen *et al.* 2008). The fact that we detected so many *trans*-eQTL may indicate that our study system has ample power to detect small effect *trans*-eQTL (our percent variance explained was 10%). Interestingly, a subset of the *trans*-eQTL we identified (located in eQTL hotspots) had exceptionally high LOD scores (75-100) that were twice as large as the largest *cis*-eQTL LOD score. Clearly, not all *trans*-eQTL have small effect sizes.

Our study demonstrates the importance of examining not just final plant height, but the developmental dynamics that contribute to height growth curves in agroecologically relevant field settings. We fit function-valued trait models to our data and, while statistically factoring out aspects of physiology such as carbon assimilation rates, demonstrate that parameters describing continuous developmental growth curves are correlated with plant fitness and yield. The shape of these growth curves (as described by *r,* d, and iD) is phenotypically plastic, while estimates of final height (*Hmax*) are relatively robust across environments. However, changes in the sign of bivariate correlations indicate a trade-off between yields at given final size vs. yields at early developmental times. We map FVT QTL to multiple chromosomes and utilize a guided eQTL mapping approach to investigate the regulatory mechanisms connecting genotype to FVT phenotype. Specifically, we use WGCNA to identify eigengenes for actin/cytoskeleton and cell division processes whose expression values that correlate with FVT traits. FVT trait seeded MR co-expression networks had an overall association with metabolic regulation and growth processes. We demonstrate that combining multiple approaches yields the best explanation of phenotypic variance. We identify more *trans*-than *cis*-eQTL and these *trans*-eQTL are highly colocalized at regulatory hotspots, likely including transcription factors that influence downstream gene regulation. Because our *cis*-and trans-eQTL hotspots colocalize with FVT QTL, these expression traits are likely components of the molecular regulatory mechanisms mediating the generation of FVT phenotypic variation from genomic variation (Fig 8).

**Fig.ure 8.**
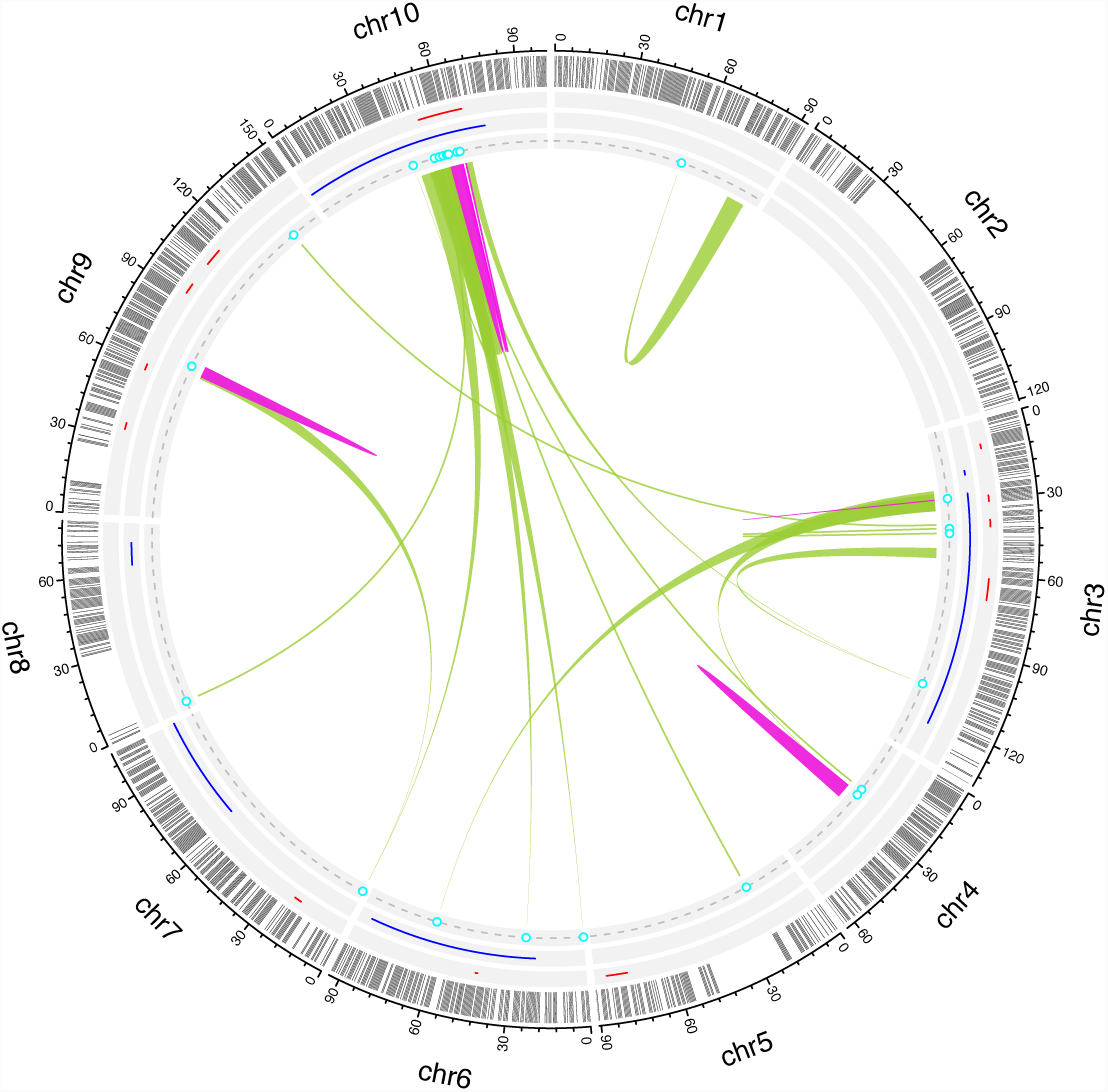
Function-Valued Trait QTL (2012 uncrowded data), Weighted Gene Co-expression Network Analysis (WGCNA) identified eigengene eQTL, and genes identified via Mutual Rank (MR) co-expression occur at regulatory hotspots on chromosomes 10 and 3, indicating that these MR genes are candidate master regulators that integrate information to generate developmental trait variation. MR gene *cis*-eQTL (pink links) on chr10 and 3 lend further credence to this relationship. MR genes with *trans*-eQTL (green links) that map to these hotspots are putative upstream genes feeding in to the FVT regulatory network. By integrating information from multiple analyses. From exterior to center: chromosomes in black, linkage map in gray, FVT QTL in red, eigengene eQTL in blue, MR genes in cyan, MR *trans*-eQTL in light green and MR *cis*-eQTL in pink.

## ACKNOWLEDGEMENTS

University of Wyoming undergraduates E. Gimpel, J. Whipps, K. Anderson, M. Pratt, J. Beckius, Blemenshine, S. Cheeney, M. Yorgason, W. Gardner, C. Planche, C. Gifford, L. Lucas, K. Riggs, Lorimer, D. Nykodym, and L. Steinken assisted with data collection and entry. C. Seals and R. Pendleton facilitated plant growth. This work is supported by National Science Foundation Grants IOS-1306574 to RLB and IOS-0923752 to CW, SW, and JM.

